# Chia (*Salvia hispanica*) gene expression atlas elucidates dynamic spatio-temporal changes associated with plant growth and development

**DOI:** 10.1101/2020.10.09.333419

**Authors:** Parul Gupta, Matthew Geniza, Sushma Naithani, Jeremy Phillips, Ebaad Haq, Pankaj Jaiswal

**Author notes:** Equal contribution and Co-First Authors (Parul Gupta, Matthew Geniza). **Correspondence:** Pankaj Jaiswal, Department of Botany and Plant Pathology, Oregon State University, Corvallis, Oregon, USA. **Abbreviations** DAF: Days after flowering; DAS: Days after sowing.

## Abstract

Chia (*Salvia hispanica* L.), now a popular superfood, is one of the richest sources of dietary nutrients such as protein, fiber, and polyunsaturated fatty acids. At present, the genomic and genetic information available in the public domain for this crop is scanty, which hinders understanding its growth and developmental processes and impedes genetic improvement through genomics-assisted methods. We report RNA-seq based comprehensive transcriptome atlas of Chia across 13 different tissue types covering vegetative and reproductive growth stages. We generated ∼394 million raw reads from transcriptome sequencing, of which ∼355 million high-quality reads were used to generate *de novo* reference transcriptome assembly and the tissue-specific transcript assemblies. After quality assessment of merged assemblies and using redundancy reduction methods, 82,663 reference transcripts were identified. Of these, 53,200 transcripts show differential expression in at least one sample and provide information on spatio-temporal modulation of gene expression in Chia. We identified genes involved in the biosynthesis of omega-3 and omega-6 polyunsaturated fatty acids, and various terpenoid compounds. The study also led to the identification of 633 differentially expressed transcription factors from 53 gene families. The coexpression analysis suggested that members of the B3, bZIP, ERF, WOX, AP2, MYB, C3H, EIL, LBD, DBB, Nin-like, and HSF transcription factor gene families play key roles in the regulation of target gene expression across various developmental stages. This study also identified 2,411 simple sequence repeat (SSRs) as potential genetic markers residing in the transcribed regions. The transcriptome atlas provides essential genomic resources for basic research, applications in plant breeding, and annotation of the Chia genome.

## Introduction

*Salvia hispanica* L. (Chia), an annual herbaceous plant originally from Central America (Cahill, 2005), is a member of the Lamiaceae (mint) family. Chia plants usually grow around one meter in height and produce raceme inflorescence bearing small purple flowers. Chia displays levels of cold and frost-tolerance, and its growth excels at higher altitudes (Ixtaina, Nolasco & Tomás, 2008; Baginsky et al., 2016). It is cultivated primarily for its nutrient-rich seeds. Chia seeds are traditionally a core component of the Mayan and Aztec population’s diet. Recently, its consumption has grown outside of South America due to its rich nutritional and gluten-free characteristics (Mohd Ali et al., 2012). The Chia seed contains approximately 40% oil by weight, of which the majority fraction is omega-3 and omega-6 polyunsaturated fatty acids (Mohd Ali et al., 2012). The seeds are gluten-free, rich in protein (15-20%), dietary fiber (20-40%), minerals (4-5%), and antioxidants (Reyes-Caudillo, Tecante & Valdivia-López, 2008; Ayerza (h) & Coates, 2009; Muñoz et al., 2013). These nutritional attributes have made Chia a desirable ‘superfood’. Several studies in humans and mouse models on a diet supplemented with Chia seed (Marcinek and Krejpcio, 2017; Oliva et al., 2013; Ullah et al., 2016; Valdivia-López and Tecante, 2015; Vuksan et al., 2017a, 2017b, 2010, 2007) report improvement in muscle lipid content, cardiovascular health, total cholesterol ratio, triglyceride content, and helped attenuate blood glucose levels in type-2 diabetes patients (Vuksan et al., 2007; Chicco et al., 2009; Peiretti & Gai, 2009; Oliva et al., 2013). Chia seeds come with variations of color and texture and may include black or dark spots. The Chia is known to show site of cultivation and environment-dependent effects on the growth of the plant, seed protein and oil content, and fatty acid composition (Ayerza, 2009). No correlation was found between nutritional composition and seed color of Chia seeds, though it is positively correlated to geographic location and environmental differences where Chia plants are grown (Ayerza (h) & Coates, 2009; Ayerza, 2010). In addition to food, Chia is a rich source of other useful products. For example, plant leaves contain various essential oil components such as β-caryophyllene, globulol, γ-muroleno, β-pinene, α-humulene, germacrene, and widdrol that are known to have insect repellant or insecticidal properties (Amato et al., 2015; Elshafie et al., 2018).

High□throughput experiments have reported large amounts of genome□wide gene expression data from various oilseed crops such as *Glycine max, Arachis hypogaea, Camelina sativa* (Libault et al., 2010; Severin et al., 2010; Clevenger et al., 2016; Kagale et al., 2016). Whereas, only a handful of studies investigated fatty acid metabolism in Chia seeds (R. V. et al., 2015; Peláez et al., 2019) and to our knowledge none across different developmental stages of Chia. As we know, plant growth and development processes are controlled by the programmed expression of a wide array of genes at Spatial and temporal scales. Gene expression atlases help predict regulatory networks and gene clusters expressed in each tissue at different developmental stages which helped in revealing the key regulators of metabolic and developmental processes (Druka et al., 2006; Sekhon et al., 2011; Stelpflug et al., 2016; Cañas et al., 2017; Kudapa et al., 2018).

In spite of Chia being one of the traditionally valued plants in South America, there are limited genetics or genomics resources available to undertake functional and comparative genomics and design plant breeding projects. Therefore, we took an initiative to build genetic and genomic resources for this important crop for the community of plant researchers and breeders. In this study, we describe tissue-specific gene expression atlas developed from 13 tissues across the vegetative and reproductive stages of Chia (Table 1). Differential expression of transcripts involved in metabolic and regulatory pathways was examined. Furthermore, we added a functional-structural annotation to transcripts and identified potential simple sequence repeat (SSRs) molecular markers and pathway enrichment to learn about important metabolic pathways. The gene expression atlas presented here is a valuable functional genomics resource and a tool for accelerating gene discovery and breeding strategies in Chia.

**Table 1:** Description of the plant material used for developing the Chia transcriptome Atlas. Samples were collected from various developmental stages and tissue types used for transcriptome analysis. DAS = Days after sowing; DAF= Days after flowering

| Growth stage | Sample collection<br>(DAS) | Sample description | Sample name |
| --- | --- | --- | --- |
| Vegetative | Day 0 | Dry Seed | Seed |
|  | Day 3 | Green cotyledon | D3-Cotyledon |
|  | Day 3 | Above ground shoot parts (whole shoot) | D3-Shoot |
|  | Day 12 | Above ground shoot parts (whole shoot) | D12-Shoot |
|  | Day 12 | Very first/youngest leaf at shoot apex | D12-P1 |
|  | Day 69 | First and Second leaves at the shoot apex | D69-P1-P2 |
|  | Day 69 | Third and fourth leaves at the shoot apex | D69-P3-P4 |
|  | Day 69 | Fifth, sixth and seventh leaves at the shoot apex | D69-P5-P6-P7 |
|  | Day 69 | Internode between 6<sup>th</sup> and 7<sup>th</sup> leaves | D69-Internode |
| Reproductive | Day 158 | Top half of the raceme inflorescence (pre-anthesis) | D158-RacemeTopHalf |
|  | Day 158 | Bottom half of the raceme inflorescence (pre-anthesis) | D158-RacemeBottomHalf |
|  | Day159 (1DAF) | Flowers from 1 day after flowering (Anthesis) | D159-Flowers |
|  | Day 164 (5DAF) | Flowers from 5 days after flowering (Anthesis) | D164-Flowers |

## Results

### Sequencing and de novo assembly

The transcriptome of Chia was generated from 13 different tissue types, including mature dry seeds, seedling shoots, leaf stages, internode, inflorescence, and flowers (Table 1). The 101 basepair (bp) length paired-end sequencing of the 39 cDNA libraries (prepared from the poly-A (mRNA) fraction of the total RNA from three biological replicates for each sample) resulted in 393,645,776 sequence reads and approximately 80Gb of the nucleotide sequence (supplementary file S1). The high-quality reads were assembled for 65 and 75 k-mer lengths, and unique transcripts were generated after merging both k-mer assemblies for each tissue type. The number of assembled transcripts were observed in the range of 27,066 to 43,491 for tissue-specific assemblies (Fig. 1A). Among vegetative tissues, D69-P1-P2 showed maximum number (43,491), and seed showed the lowest number (27,066) of assembled transcripts (Fig. 1A). Among reproductive tissues, the maximum number of transcripts (43,418) with the highest average length of about 1000 bases was observed in the top half part of the D158-Raceme inflorescence (Fig. 1A). Total high-quality paired-end reads (352,976,255) from all tissue libraries were pooled and assembled at 67 and 71 k-mer lengths using Velvet (Zerbino & Birney, 2008) and Oases (Schulz et al., 2012). Chia transcript isoforms generated by each k-mer (67 and 71 k-mer lengths) assembly were consolidated (referred to as merged assembly) to represent the total number of 145,503 unique transcripts of ≥201 bases in length (Fig. 1B).

**Figure 1:**
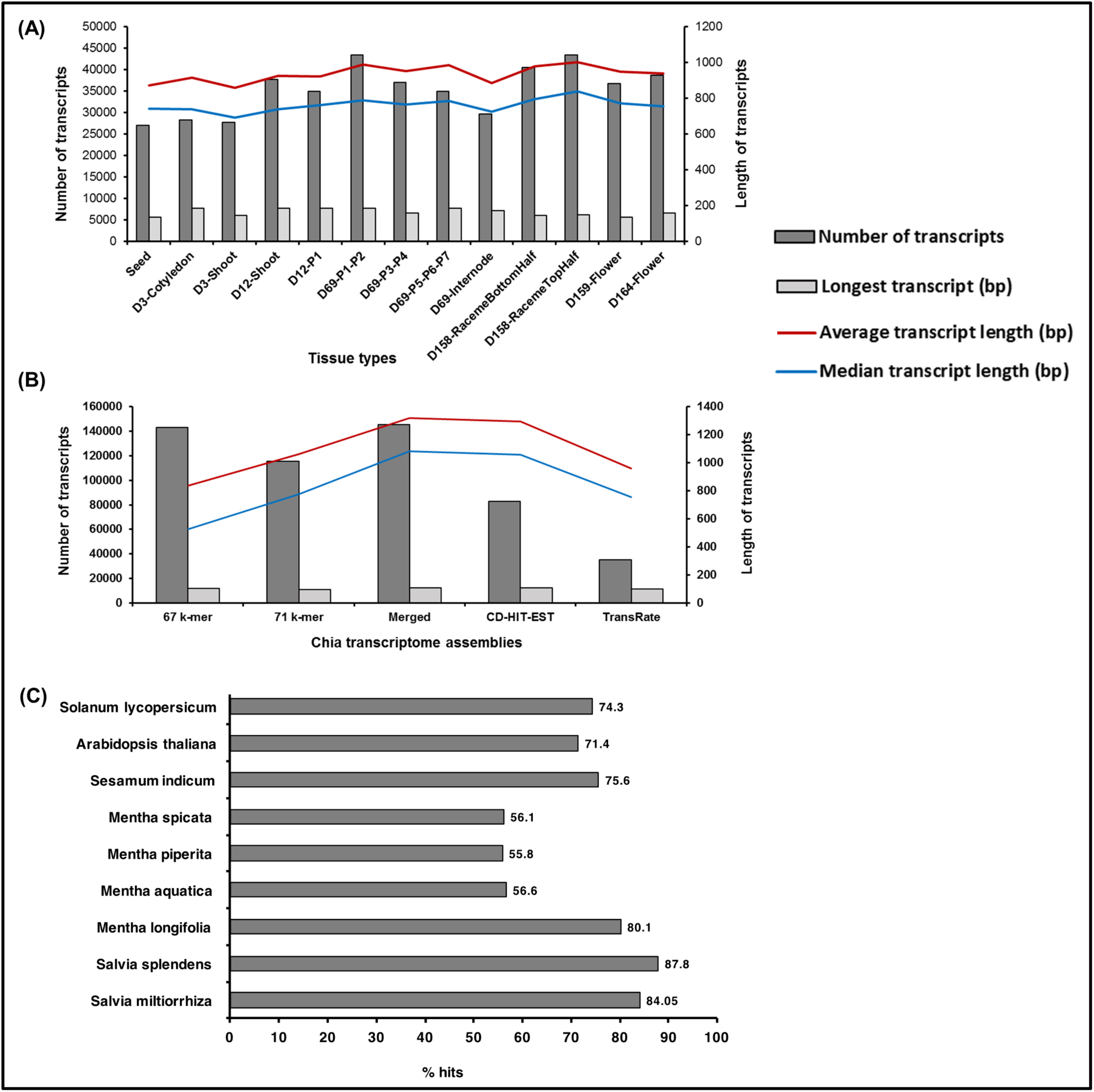
Statistics of *S. hispanica* transcriptome assemblies and BLAST results. **(A)** tissue-specific assembly; **(B)** reads from each tissue types are combined and assembled at 67 k-mer and 71 k-mer, Merged assembly of 67 k-mer and 71 k-mer, CD-HIT-EST and TransRate assemblies by removing redundant reads; **(C)** Comparison of *S. hispanica* transcripts with publically available Lamiales and eudicot gene models and peptide set.

As part of the quality assessment addressing redundancy, we first used the CD-HIT-EST algorithm (Li & Godzik, 2006) to reduce the number of redundantly assembled transcripts by grouped sequences displaying similarities higher than 90%. This yielded 82,663 transcripts (Fig. 1B). This step was followed by running the transcriptome quality assessment software, TransRate (Smith-Unna et al., 2016). TransRate detects the redundant transcripts by aligning the reads to multiple transcripts, but the assignment process assigns them all to the transcript that best represents the canonical form. This process reduced the originally assembled transcriptome (145,503 transcripts) to 35,461 transcripts (Fig. 1B). We observed that the assembly produced by CD-HIT-EST experienced little to no loss in percentage of reads aligned. The assembly produced by TransRate, which utilizes Salmon (Patro et al., 2017) to estimate transcript abundance using map-based methods, contained nearly 50% less reads aligned in comparison to the CD-HIT-EST assembly. Furthermore, we used quality assessment tool QUAST (Mikheenko et al., 2016) on the original assembly and each of the redundancy reduced assemblies (Supplementary file S2). The original and TransRate assemblies both had the better statistics in transcript number and length and both assemblies also contained the worst statistics in the complementing category (Supplementary file S2). The assembly produced by CD-HIT-EST represented the most moderate version of the assembly. Using the quality assessment and alignment data as criteria, we decided that the CD-HIT-EST assembly with 82,663 transcripts would be the most appropriate for downstream analyses. Workflow for assembly and downstream analysis is showed in Supplementary file S3.

### Functional annotation of Chia transcriptome

We compared the 82,663 assembled Chia transcripts to publicaly available genomes and gene models of Eudicots using BLASTx and tBLASTx (Mount, 2007) to estimate approximate coverage of genes represented in the assembled transcriptome (Fig.1C). More than 84% of assembled Chia transcripts mapped to the closely related *Salvia miltiorrhiza* (Wenping et al., 2011) and *Salvia splendens* (Ge et al., 2014) transcriptomes (Fig. 1C). The dispersion of coverage within the genus is not surprising since the *Salvia* genus is very diverse. Both *S. miltiorrhiza* and *S. splendens* share a common center of origin in China, whereas *Salvia hispanica* originated in Central America. Within the Lamiaceae, about 56% of the transcripts mapped to members of the *Mentha* (mint) genus, namely, Watermint (*M. aquatica*), Peppermint (*M. piperita*), and Spearmint (*M. spicata*) (Ahkami et al., 2015a). Moving up the taxonomic rank to the order of Lamiales, 75% of Chia transcripts mapped to sesame (*Sesamum indicum*) (Zhang et al., 2013), an oilseed crop. A total of 71% and 74% of assembled Chia transcripts aligned to the model plant *Arabidopsis thaliana* and the *Solanum lycopersicum* (tomato) proteome set, respectively (Fig. 1C). Although assembled transcriptomes were not available, the RNA-Seq reads from two recently sequenced and publicly available *Salvia hispanica* projects (Sreedhar et al., 2015; Boachon et al., 2018) for seed (INSDC Accession PRJNA196477) and leaf tissues (INSDC Accession PRJNA359989) were aligned against our assembled chia transcriptome. About 69% sequence reads from the seed, and 43% of the leaf transcriptome sequences mapped to our assemblies.

Peptide sequences from the assembled transcripts were generated using TransDecoder, which scans all ORFs based on homology searches from Pfam and BlastP as ORF retention criteria. Out of total 82,663 transcripts, 65,587 transcripts from Chia were translated into 99,307 peptides. The number of peptides is higher than the number of transcripts assembled due to multiple open reading frames (ORFs) occurring in a single transcript. Functional annotation of peptides was first carried out using InterProScan (Jones et al., 2014a) to assign structural-functional domains and then by employing agriGO (Du et al., 2010b). We were successful in assigning InterPro accessions to the 45,209 peptides (Supplementary file S4) and Gene Ontology (GO) terms to a total of 32,638 peptides (Supplementary file S5). A total of 20,857 peptides were with GO biological process (BP); 8,677 peptides were associated to GO cellular component (CC), and 26,877 peptides were annotated to GO molecular function (MF) terms (Supplementary file S5).

### Development of gene expression atlas

A final set of 82,663 assembled transcripts and the RSEM (Li & Dewey, 2011b) package was used to estimate transcript abundance based on FPKM (Fragments Per Kilobase of transcript per Million mapped reads). After removing transcripts with extremely low/insignificant expression, we considered 82,385 transcripts for further analysis. In order to visualize cross-sample comparison, a heatmap of distance matrix was generated that showed hierarchical clustering of Pearson’s correlations based upon FPKM values for all transcripts (Fig. 2). Most of the tissues clustered together based on developmental attributes that provide an intriguing clue about the spatial and temporal scale of the samples pattern. For example, vegetative tissues, D3 (cotyledon and shoot) and D12 (shoot and very first leaf at shoot apex), clustered together. Leaf stages varied at maturity level were also clustered together. Interestingly, we observed that seed and internode tissues clustered together, suggesting that they share common transcripts. Similarly, among reproductive tissues, flowers (D159 and D164) and inflorescence tissues (raceme top and bottom half) clustered together.

**Figure 2:**
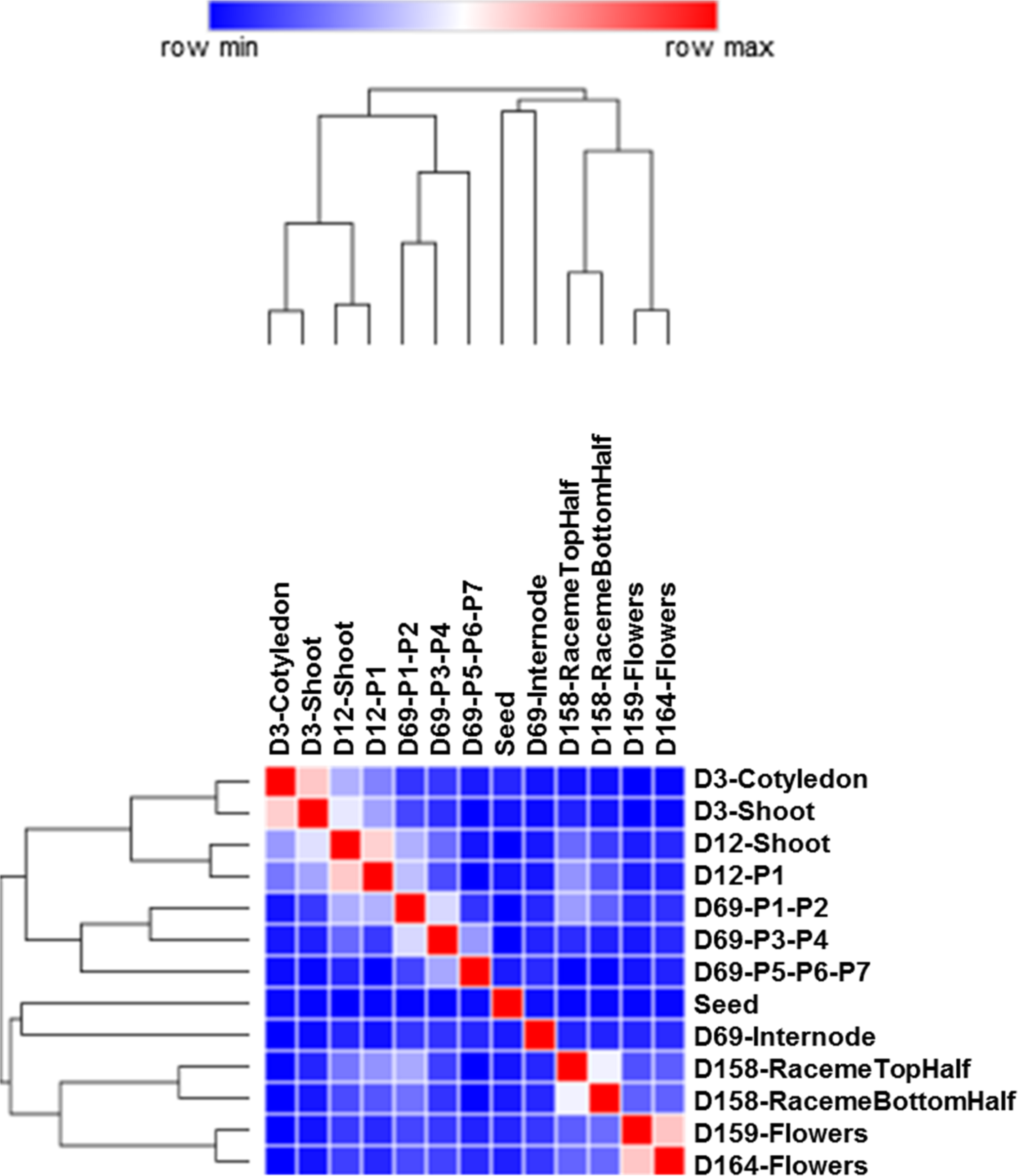
Gene expression patterns across different tissues of Chia. Heatmap of hierarchical clustering of the Pearson correlations for all 13 tissues included in the gene expression atlas. Log2 transformed FPKM values were used for the similarity matrix of transcripts. The color scale indicates the degree of correlation.

In order to study the gene clusters with a similar expression, the expression trend of all transcripts across developmental stages were represented in 20 clusters (Supplementary file S6). Most of the transcripts in cluster #1 (7507 transcripts), #5 (4616 transcripts), #12 (3909 transcripts), #15 (3619 transcripts), #18 (2679 transcripts), and #20 (2173 transcripts) showed higher expression in seeds, D3-tissues (cotyledon and shoot), mature leaf stage (D69-P5-6-7), flowers (D-159 and D-164), inflorescence (top half and bottom half) and internode, respectively (Supplementary file S6). Transcripts in cluster #1 enriched for LEA (Late embryogenesis abundant), seed storage proteins, oil body-associated proteins, and oleosin family members (Supplementary file S7). In soybean seed transcriptomes, storage protein genes like beta-conglycinins, oleosins, glycinins, several LEA proteins and dehydrin genes showed higher expression with respect to other genes (Severin et al., 2010; Jones & Vodkin, 2013). Cluster #5 was considered rich in transcripts required for initial growth (D3-Cotyledon, D3-Shoot) of seedling after germination. The majority of highly expressed transcripts were annotated as zinc finger, basic leucine zipper family members, photosystem I and II related proteins, aquaporins, and calcineurin-like phosphoesterase domain-containing proteins (Supplementary file S7). In cluster #12, highly expressed transcripts in D69-P5-6-7 leaf stage were annotated as disease resistance proteins, leucine-rich receptor kinases (LRR-RLKs), and wall-associated receptor kinases (WAKs) (Supplementary file S7). Transcripts that encode transporter (ABC, phosphate, aluminum transporters) proteins, cytochrome P450s, glycosyltransferases, and WRKY transcription factors also enriched in this cluster. Cluster 15 represents transcripts that showed higher expression in flowers. Transcripts annotated as beta-glucosidase, multidrug and toxic compound extrusion proteins, cinnamyl alcohol dehydrogenase (involved in lignin biosynthesis in floral stem in *Arabidopsis*) (Sibout et al., 2005), germin-like proteins (might play a role in plant defense), pectin acylesterases, MYB family transcription factors (MYB21 and MYB24), GDSL lipase family members, and cytochrome P450s were highly enriched in this cluster (Supplementary file S7). MYB21 and MYB24 transcription factors are known for their role in petal, stamen, and gynoecium development in flowers (Reeves et al., 2012). Cinnamyl alcohol dehydrogenases are involved in lignin biosynthesis in floral stem in Arabidopsis (Sibout et al., 2005), and germin-like proteins play an important role in response to pathogens (Zimmermann et al., 2006; Manosalva et al., 2009; Wang et al., 2013). Transcripts that are highly upregulated in inflorescence tissues grouped in cluster #18. Transcription factors that play a vital role in floral meristem development enriched in this cluster. For example - agamous-like MADS-box proteins and MYB family transcription factors (Supplementary file S7). MYBs and MADS-box transcription factors are essential regulators of various developmental processes (Zimmermann et al., 2004; Millar & Gubler, 2005; Yang et al., 2007; Gomez et al., 2011; Kobayashi et al., 2015). Cluster #20 enriched with the transcripts upregulated in the D69-Internode sample (Supplementary file S7). It includes expression of transcription factors from the– MYB (MYB54, MYB52) and NAC domain-containing transcription factor families known for their role in the development of the vegetative internodes. MYB54, MYB52, and NAC transcription factors are also known to regulate secondary cell wall biosynthesis (Zhong et al., 2008; Grant et al., 2010; Cassan-Wang et al., 2013). Transcripts encode xyloglucan endotransglucosylase, which participates in cell wall construction of growing tissues, were also upregulated in internode (cluster 20) compared to other tissues. A set of transcripts encode for receptor-like protein kinases, involved in the signaling pathways known to regulate cell expansion (Guo et al., 2009; Haruta et al., 2014) is upregulated in cluster 20.

### Differential expression at each growth stage

A total of 53,200 unique transcripts were differentially expressed among all tissue types of which 38,480 transcripts show log_2_ fold change ≥ 2. Seed shows the highest number of differentially expressed transcripts, followed by D69-P5-P6-P7, D69-Internode, and D12-P1 (Table 2). Only D3-cotyledon showed the higher number of transcripts were under the upregulated category compared to the downregulated ones, whereas in the other 12 tissues, this pattern was opposite (Table 2). Seed showed the maximum number of tissue-specific differentially expressed transcripts (13,450) followed by D69-P5-P6-P7, D69-internode, D12-P1, and D3-Cotyledon tissue types (Table 2). The maximum number of upregulated transcripts was observed in seed (6,284) followed by D3-Cotyledon (2,632), D69-P5-P6-P7 (1,884), D69-Internode (1,390), and D159-Flowers (1,274). Similarly, the maximum number of downregulated transcripts were also observed in seed (13,429), followed by D69-P5-P6-P7 (6,637), D69-Internode (5,163), D12-P1 (3,976), and D164-Flowers (3,353).

**Table 2:** Differentially expressed (DE) transcripts across various developmental stages

| Tissue type | Total DE | Tissue specific | Upregulated<br>(log<sub>2</sub> FC ≥ 2) | Downregulated<br>(log<sub>2</sub> FC ≤ -2) |
| --- | --- | --- | --- | --- |
| Seed | 28,641 | 13,450 | 6,284 | 13,429 |
| D3-Cotyledon | 7,377 | 1,781 | 2,632 | 1,746 |
| D3-Shoot | 3,415 | 495 | 970 | 1,161 |
| D12-Shoot | 2,136 | 270 | 52 | 1,521 |
| D12-P1 | 8,795 | 2,038 | 770 | 3,976 |
| D69-P1-P2 | 3,511 | 633 | 288 | 2,185 |
| D69-P3-P4 | 3,019 | 556 | 479 | 1,701 |
| D69-P5-P6-P7 | 14,140 | 3,504 | 1,884 | 6,637 |
| D69-Internode | 9,260 | 2,152 | 1,390 | 5,163 |
| D158-RacemeTopHalf | 5,591 | 1,183 | 770 | 2,865 |
| D158-RacemeBottomHalf | 3,614 | 804 | 852 | 1,860 |
| D159-Flowers | 6,047 | 1,136 | 1,274 | 2,883 |
| D164-Flowers | 6,134 | 879 | 969 | 3,353 |

Besides, the distribution of differentially expressed transcripts between different combinations of similar or related developmental stages was evaluated (Fig. 3). In the initial growth stages: seed, D3-cotyledon, D3-shoot, and D12-shoot tissues, only 213 differentially expressed transcripts were common, and 70%, 8.9%, 2.4%, and 2.5% transcripts were specific to seed, D3-Cotyledon, D3-shoot and D12-shoot tissues, respectively (Fig. 3A). The majority of transcripts highly upregulated (log_2_ fold change ≥ 10) in seed but downregulated (log_2_ fold change ≤ −4) in other initial growth stages are seed storage, and LEA proteins. Among all leaf developmental stages, only 1.6% of differentially expressed transcripts were common, and 5372, 1599, 1092, and 9938 transcripts were specific to D12-P1, D69-P1-P2, D69-P3-P4, and D69-P5-P6-P7, respectively (Fig. 3B). Transcripts encoding for LRR-RLKs, WAKs, and RHD3 domain-containing proteins were highly expressed in D69-P5-6-7 compared to other leaf developmental stages. In early leaf developmental stages (D12-P1 and D69-P1-P2), transcripts encoding for Growth Regulating Factors (GRF2, GRF5) and bHLH domain-containing (SPEECHLESS) transcription factors were highly expressed compared to those in the mature leaf stages. GRF transcription factors play an important role in leaf growth, and the bHLH SPEECHLESS factors are involved in stomata initiation and development (Kim et al., 2003; MacAlister et al., 2007; Kanaoka et al., 2008; Lampard et al., 2008). Among raceme inflorescence and flower tissues, only 2.8% (414) of differentially expressed transcripts were common, and 3315, 1689, 2591, and 2648 transcripts were specific to RacemeTopHalf, RacemeBottomHalf, D159-Flowers, and D164-Flowers, respectively (Fig. 3C). The higher expression of transcripts that encode ZFP2 and MYB transcription factors, cinnamyl alcohol dehydrogenase, and pectin acylesterases showed up in flowers. ZFP2 controls floral organ abscission (Cai & Lashbrook, 2008), and cinnamyl alcohol dehydrogenases are involved in lignin biosynthesis in floral stem (Sibout et al., 2005). Transcripts annotated as terpene synthases show upregulation in flowers as compared to the inflorescence tissues. Transcripts annotated as oxidation-reduction related activities were highly enriched in flowers, inflorescence, D-69 leaf stages, and internode tissues, which indicated that ROS concentration increases during these growth stages as in other species (Rogers, 2012; Rogers & Munné-Bosch, 2016; Singh et al., 2016).

**Figure 3:**
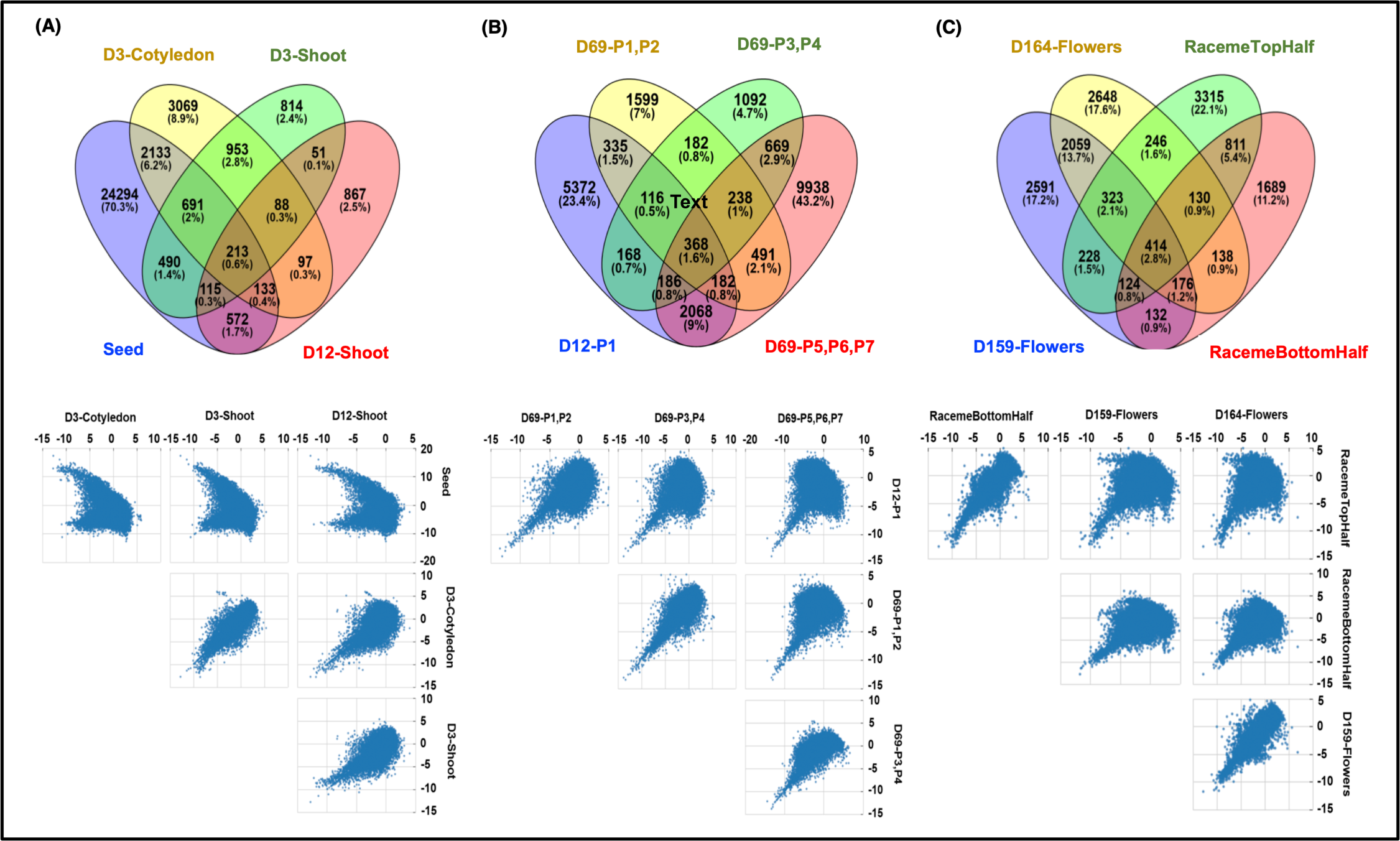
Differential expression of Chia transcripts among **(A)** seed, D3-cotyledon, D3-shoot, and D12-shoot; **(B)** D12-P1 and D69-P1-P2, D69-P3-P4, D69-P5-P6-P7; **(C)** reproductive stages including RacemeTop and BottomHalf tissues, D159- and D164-flowers. Vein diagrams in the upper panel represent common and unique differentially expressed transcripts in each tissue type, and scatter plots in the lower panel represent the distribution pattern of differentially expressed transcripts across each tissue type.

### Pathways enriched across different development stages

The metabolic network representation across developmental stages of Chia were determined by mapping to KEGG (Kyoto Encyclopedia of Genes and Genomes) pathways. A total of 5,555 transcripts mapped to 464 pathways. The higher numbers of transcripts mapped to starch and sucrose metabolism (PATH:ko00500), fatty acid metabolism (PATH:ko01040), phenylpropanoid biosynthesis (PATH:ko00940), photosynthesis (PATH:ko00195), fatty acid biosynthesis (PATH:ko00061), and various amino acids metabolism processes (Supplementary file S8). The expression pattern of transcripts encoding the enzymes for fatty acid metabolism and unsaturated fatty acid (including omega-3 and omega-6) metabolism across different developmental stages was analyzed(Figure 4). Transcripts for acetyl-CoA carboxylase (EC 6.4.1.2), the very first enzyme catalyzing the conversion of acetyl-CoA to malonyl-CoA in the fatty acid biosynthesis were highly expressed in all tissues except seeds. The malonyl group from malonyl-CoA is transferred to acyl carrier proteins (ACP) in the next step for further elongation. We identified transcripts for all the enzymes participating in the elongation steps. Acyl-ACP thioesterases (3.1.2.14) acts in the last steps of fatty acid biosynthesis and serves as a determining factor for the generation of a variety of fatty acids within an organism. Since, Chia seeds are very rich in unsaturated fatty acids: linoleic and α-linolenic acids, genes involved in unsaturated fatty acids biosynthesis were queried for their expression pattern across all tissue types (Fig. 4B). Fatty acid desaturases (FADs) are the crucial enzymes to perform the desaturation of fatty acids. We identified 32 FAD transcripts from FAD2, FAD3, FAD6, FAD7and FAD8 families (Table 3). Endoplasmic reticulum localized *FAD2* and plastid localized *FAD6* encode two ω-6 desaturases required to convert oleic acid to linoleic acid (18:2^Δ9,12^) (Zhang et al., 2012). The desaturation of linoleic acid (18:2^Δ9,12^) to α-linolenic acid (18:3^Δ9,12,15^) is catalyzed by the endoplasmic reticulum localized FAD3 and plastid localized FAD7 and FAD8 proteins (Dar et al., 2017; Xue et al., 2018).

**Table 3:** Transcripts annotated as fatty acid desaturase

| **Fatty acid desaturases** | **Transcripts** |
| --- | --- |
| FAD2 | Sh_Salba_v2_49454<br>Sh_Salba_v2_49451<br>Sh_Salba_v2_66763 |
| FAD3 | Sh_Salba_v2_74023<br>Sh_Salba_v2_93044<br>Sh_Salba_v2_93043<br>Sh_Salba_v2_74025<br>Sh_Salba_v2_93046<br>Sh_Salba_v2_74024<br>Sh_Salba_v2_74022<br>Sh_Salba_v2_93047<br>Sh_Salba_v2_93045 |
| FAD6 | Sh_Salba_v2_05727<br>Sh_Salba_v2_05731<br>Sh_Salba_v2_05730<br>Sh_Salba_v2_05725<br>Sh_Salba_v2_05728<br>Sh_Salba_v2_05721<br>Sh_Salba_v2_05724 |
| FAD7 | Sh_Salba_v2_69172 |
| FAD8 | Sh_Salba_v2_69173<br>Sh_Salba_v2_90850<br>Sh_Salba_v2_52578<br>Sh_Salba_v2_69162<br>Sh_Salba_v2_52570<br>Sh_Salba_v2_52575<br>Sh_Salba_v2_52576<br>Sh_Salba_v2_69169<br>Sh_Salba_v2_69166<br>Sh_Salba_v2_52573<br>Sh_Salba_v2_69174<br>Sh_Salba_v2_69171 |

**Figure 4:**
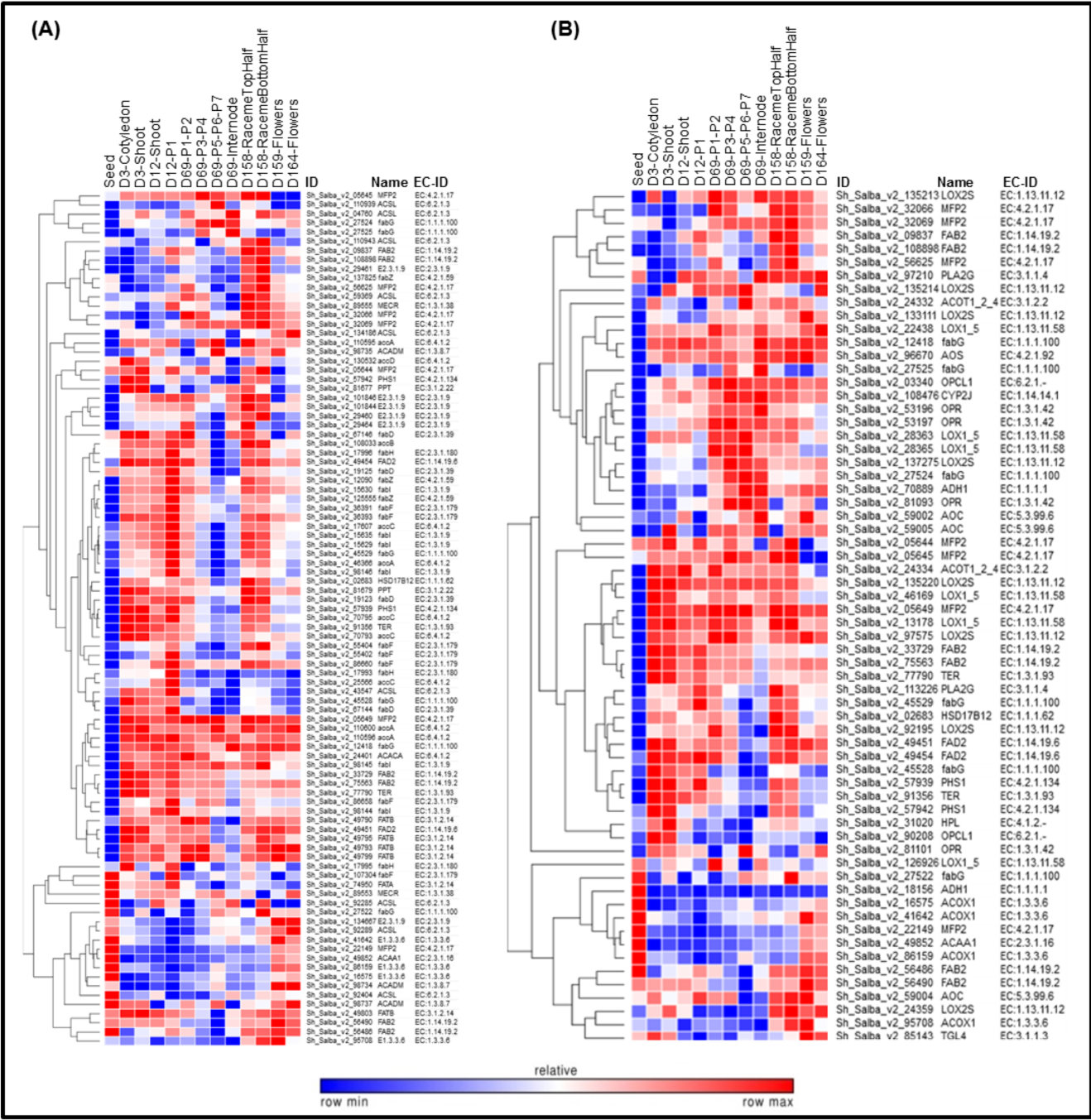
Expression pattern of transcripts involved in fatty acid metabolism across tissue types. **(A)** Fatty acid metabolism; **(B)** Unsaturated fatty acids, Omega-3 (*α*-Linolenic acid) and Omega-6 (Linoleic acid) fatty acids metabolism

Mint, a Lamiaceae family plant, is primarily known for the production of monoterpenes, e.g., menthol and limonene (Aharoni, Jongsma & Bouwmeester, 2005; Ahkami et al., 2015b); however, the majority of Chia terpenes are sesqui-, di-, and tri-terpenes (Ma et al., 2012; Trikka et al., 2015; Cui et al., 2015). In our Chia dataset, we observed the expression profile of transcripts involved in the biosynthesis of terpenoid backbone, monoterpenes, and sesquiterpenes. Transcripts encoding enzymes for each catalytic step of terpenoid backbone synthesized by the MEP (2-C-methyl-D-erythritol 4-phosphate) and the mevalonate (MVA) pathways showed differential expression pattern among all tissue types (Fig. 5). Transcripts for monoterpene synthases such as 1,8-cineole synthase (EC 4.2.3.108), myrcene synthase (EC 4.2.3.15), and linalool synthase (EC 4.2.3.25) were highly expressed in reproductive tissues (Fig. 5), indicating that flowers are the prime site for the biosynthesis of essential oils known to have therapeutic properties. However, transcripts for the sesquiterpene synthases, β-caryophyllene synthase (EC 4.2.3.57), α-humulene synthase (EC 4.2.3.104), Germacrene synthase (EC 4.2.3.60), and solavetivone oxygenase (EC 4.2.3.21), known for plant herbivory defense enriched in the vegetative tissues (Fig. 5).

**Figure 5:**
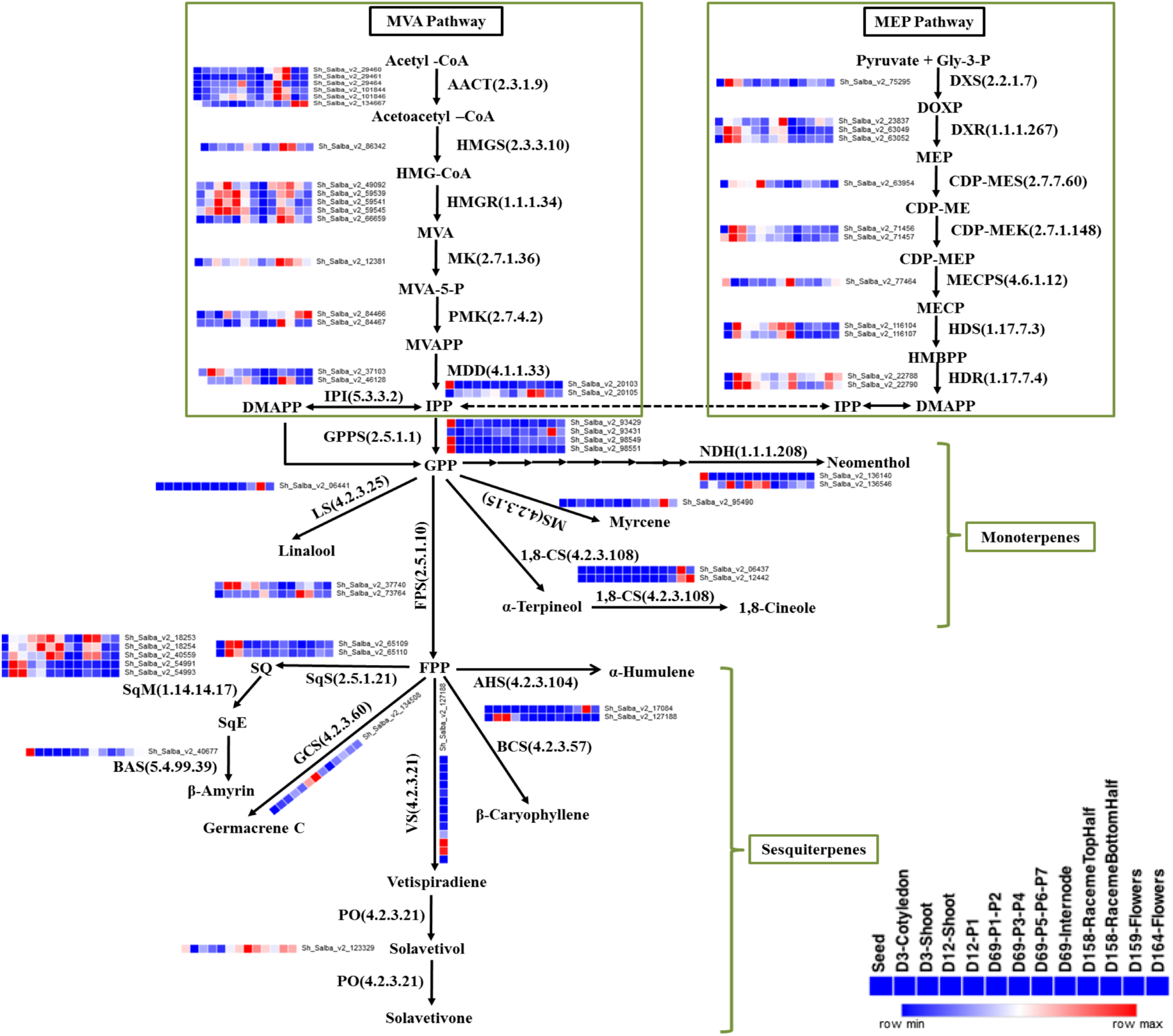
Expression pattern of transcripts involved in biosynthesis of terpenes across tissue types. Biosynthesis of IPP, a central precursor for other terpenes biosynthesis, via cytosolic MVA (mevalonate) and plastid localized MEP (2-C-methyl-D-erythritol 4-phosphate) pathways. Biosynthesis of various monoterpenes from GPP and sesquiterpenes from FPP. AACT, Acetyl-CoA acetyltransferase; HMG-CoA, 3-hydroxy-3-methylglutaryl-CoA; MVA, mevalonate; MVA-5-P, mevalonate 5-phosphate; MVAPP, mevalonate diphosphate; IPP, isopentenyl diphosphate; DMAPP, dimethylallyl diphosphate; GPP, geranyl diphosphate; FPP, farnesyl diphosphate; HMGS, HMG synthase; HMGR, HMG reductase; MK, mevalonate kinase; PMK, phosphomevalonate kinase; MDD, Mevalonate diphosphosphate decarboxylase; IPI, IPP isomerase; GPPS, geranyl diphosphate synthase; FPPS, FPP synthase; Gly-3-P, glyceraldehyde-3-phosphate; DOXP, 1-deoxy-D-xylulose-5-phosphate; MEP, 2-C-methyl-D-erythritol-4-phosphate; CDP-ME, 4-diphosphocytidyl-2-C-methyl-D-erythritol; CDP-MEP, 4-diphosphocytidyl-2-C-methyl-D-erythritol-2-phosphate; MECP, C-methyl-D-erythritol-2,4-diphosphate; HMBPP, hydroxy methylbutenyl-4-diphosphate; DXS, DOXP synthase; DXR, DOXP reductoisomerase; CDP-MES, 2-C-methyl-D-erythritol4-phosphatecytidyl transferase; CDP-MEK, 4-(cytidine-5-diphospho)-2-C-methyl-D-erythritol kinase; MECPS, 2,4-C-methyl-D-erythritol cyclodiphosphate synthase; HDS, 1-hydroxy-2-methyl-2-(E)-butenyl-4-phosphatesynthase; HDR, 1-hydroxy-2-methyl-2-(E)-butenyl-4-phosphate reductase; NDH, neomenthol dehydrogenase; MS, myrcene synthase; 1,8-CS 1,8-cineole synthase; LS, linalool synthase; AHS, alpha-humulene synthase; BCS, beta-caryophyllene synthase; VS, vetispiradiene synthase; PO, premnaspirodiene oxygenase; SQ, squalene, SqS, squalene synthase; SqE, squalene epoxide; SqM, squalene monooxygenase; BAS, beta-amyrin synthase; GCS, germacrene C synthase.

### Transcription factor network

Transcription factors are the key regulators that control many biological processes in plants, including growth and development. To gain detailed information about transcription factors, we investigated Chia transcriptome and identified 633 differentially expressed transcripts annotated to 53 transcription factor families (Supplementary file S9). The highest number of transcripts belong to MYB (60), followed by bHLH (45), NAC (38), bZIP (32), WRKY (28), C2H2 (27), MYB-related (25), MADS-box (26), C3H (24), G2-like (22), Hd-ZIP (22), Trihelix (17), TCP (14), Dof (13), GATA (13), GRAS (13), and TALE (13) gene families, etc.. The expression pattern of differentially expressed transcription factors across developmental stages is shown in Fig. 6A.

**Figure 6:**
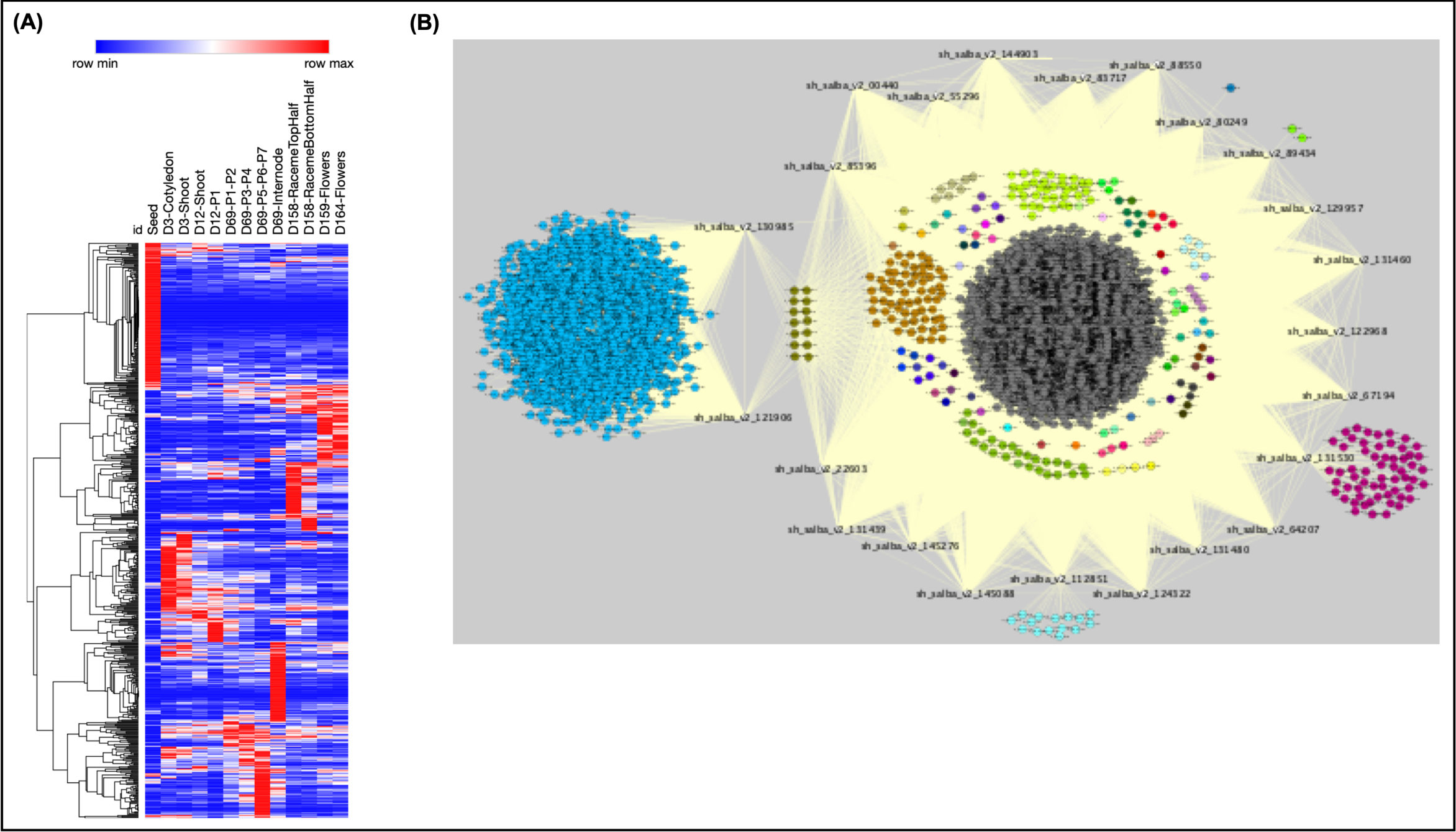
Expression pattern of transcription factors and coexpression analysis with differentially expressed transcripts. **(A)** Expression pattern of differentially expressed transcription factors (633) across various developmental stages. **(B)** Coexpression network of 23 highly upregulated (log_2_ fold change ≥ 5) transcription factors (bait) and differentially expressed transcripts (11,055) with log_2_ fold change ≥ 2. Bait transcripts are shown in white color nodes with corresponding transcripts IDs, whereas correlated transcripts are represented as colored nodes. Each set of nodes is represented with different colors based on the number of correlating edges (yellow lines) connected to that node. For example-In a set of blue color nodes (1593), each transcript (blue node) showing 2 edges connected to two bait transcripts (Sh_Salba_v2_130985, Sh_Salba_v2_121906).

To gain insight into the regulatory role of transcription factors in Chia, we filtered out highly upregulated transcription factors in any of the 13 tissues (≥ 5 log_2_ fold change) to build a coexpression network. In an in-silico experiment, we used 23 transcription factors as baits (nodes) and FPKM values of 38,480 transcripts as an expression matrix. This analysis revealed a total of 1,98,746 connections (edges) among 23 bait transcript nodes and 11,055 differentially expressed transcript nodes (Fig. 6B). Two transcription factors, Sh_Salba_v2_130985, Sh_Salba_v2_121906 highly expressed in D69-Internode but downregulated or absent in other tissues, were annotated as MYB and C3H family members, respectively. Both the transcription factors connected to a set of 1,593 transcripts that showed no connection to any other bait nodes. A set of 16 transcripts solely connected to sh_salba_v2_112851 an ERF transcription factor that was highly upregulated (log_2_ fold change 5.561) in seed. All correlated 15 transcripts were downregulated in seed and other tissue types. The MYB transcription factor transcript sh_salba_v2_131530 was upregulated in seed and connected to a set of 59 transcripts that were downregulated in seed and other tissues. Two transcripts (sh_salba_v2_32610, sh_salba_v2_03332), that downregulated in seed showed a connection to B3-domain containing sh_salba_v2_89434 bait. A transcript (sh_salba_v2_86132), downregulated in seed and annotated as disease resistance protein correlated to a HSF transcription factor bait transcript sh_salba_v2_80249. A set of 14 transcripts downregulated in seed and D69-Internode were connected to all 23 bait transcripts. Bait transcripts also correlated to each other suggesting a multiple regulatory modules within the network (Fig. 6B).

### Identification of Simple Sequence Repeat molecular markers

Simple Sequence Repeats (SSRs) are an important class of genetic markers widely used in molecular breeding applications. SSRs identified from the transcriptome are highly advantageous as compared to SSRs identified from the genome. If the SSRs identified from the transcribed region is polymorphic, they may have a direct impact on the expression, structure, stability of the open reading frame, and altered peptide sequence and functional domains. We identified a total of 2,411 SSRs in the *de novo* assembled transcriptome represented by di-, tri-, and tetra-nucleotide motifs (Supplementary file S10). The most abundant di-, tri, and tetra-nucleotide motifs were CT (201), GAA (84), and AGTC (12), respectively (Supplementary file S11). A total of 1,771 SSRs were present in the significantly differentially expressed transcripts, and 148 SSR markers found in the expressed transcripts mapped to at least one metabolic pathway (Supplementary file S12).

## Discussion

At present, the genetic information and genomic resources on Chia are scanty. Before this work, a couple of studies focused on the expression of lipid biosynthesis genes in developing Chia seeds has been reported (Sreedhar et al., 2015; Peláez et al., 2019; Wimberley et al., 2020). Big data biology can fill in this gap and build reference resources for breeding and improvement of this important crop. Using RNA-Seq coupled with the *de-novo* transcriptome assembly approach, we developed a comprehensive gene-expression atlas for Chia from 13 different tissue samples (see Table 1) collected at various developmental stages of plants. Assembled transcripts were annotated using BLASTx and tBLASTx and then translated into peptides using Transdecoder (v2.1.0) with a minimum peptide length of 50 or more amino acids. The derived peptide set was subjected to InterProScan (Zdobnov & Apweiler, 2001a) and AgriGO (Du et al., 2010a) analyses to assign structurally conserved domains and GO terms. Overall, the Chia transcriptome dataset is diverse, representing a majority of peptides belong to the cellular metabolic process, catalytic activity, regulation of gene expression, transport, ion binding, organelle, nucleus and macromolecular complexes. A comparison of Chia transcripts data sets to genomic/transcriptome datasets (Figure 1C) from the six most closely related eudicots including topmost matching with transcripts of perennial herbs, the red sage *Salvia miltiorrhiza* (Wenping et al., 2011) and the scarlet sage, *Salvia splendens* (Ge et al., 2014) - both species-rich in secondary metabolites known for their ue in traditional medicine. In *de novo* assembled transcripts, the read mapping ambiguity is prevalent, and other popular tools, such as edgeR (Robinson, McCarthy & Smyth, 2010) and DESeq (Anders & Huber, 2010) do not take variance due to read mapping uncertainty into consideration. Therefore, we employed EBSeq (Leng et al., 2013) for conducting differential gene expression analysis that takes variance due to the sequence read mapping ambiguity into account by grouping the isoforms.

This comprehensive expression atlas facilitated in the mining of gene expression data for regulatory and metabolic processes, tissue-specific gene expression pattern, and provided insights about functional relatedness of genes and their expression across developmental stages. Hierarchical clustering of Chia transcripts suggested the role of different gene families in the development of each growth stage, thus providing a foundation for studying the molecular mechanisms occurring in different tissues and developmental stages. For example, seed-specific transcripts: seeds are rich in storage, and LEA proteins are required for seed germination and embryogenesis. The Leaf-specific transcripts: mature leaves have higher expression of LRR-RLKs and WAKs proteins. LRR-RLKs are involved in guard cells and stomatal patterning (Shpak et al., 2005), and resistance to pathogens. GRF family transcription factors play an essential role in the growth and development of leaf, were highly expressed in D69-P1-P2 leaf stages. In Arabidopsis, GRF1, GRF2, and GRF5 regulate leaf number and size (Kim, Choi & Kende, 2003; Horiguchi, Kim & Tsukaya, 2005; Lee et al., 2009).

The flower-specific transcripts show higher expression of terpene synthases, which suggested that as a characteristics of Lamiaceae family, Chia flowers are also rich in monoterpene synthases, e.g., 1, 8-cineole synthase (EC 4.2.3.108) and β-myrcene synthase (EC 4.2.3.15). Cineole and myrcene are found in fragrant plants and are known to have therapeutic properties such as sedative, anti-inflammatory, antispasmodic, and antioxidant (do Vale et al., 2002; Moss & Oliver, 2012; Bouajaj et al., 2013; Juergens, 2014; Khedher et al., 2017). The reproductive versus vegetative tissue comparison shows that monoterpene synthases were expressed highly in reproductive tissues, and sesquiterpene synthases were prominent in vegetative tissues. These findings confirm that flowers are involved in the synthesis of fragrance and therapeutic essential oils, whereas vegetative tissues are rich in herbivory defense and insecticidal compounds.

Chia seeds are a rich source of polyunsaturated fatty acids. We observed lower expression of FAD transcripts in seeds as compared to other tissue types. This suggested that seed might serve as a storage organ for polyunsaturated fatty acids rather synthesis site or seeds we used in this study were dry and in semi-dormant condition. Essential oils, the secondary metabolic plant products of the terpenoid pathway produced by Lamiaceae plant family members, are highly desired for their usage in medicine, food, cosmetics, and for their agronomic properties such as insecticides, herbivory, and pathogen defense. In this Chia dataset, we identified transcripts encoding enzymes for terpenoid backbone (MVA and MEP) pathways. Monoterpene synthases are involved in essential oil biosynthesis, and sesquiterpene synthases are primarily involved in the biosynthesis of insecticidal compounds. Phenylpropanoid and flavonoid biosynthesis pathways are also highly enriched in seeds and other tissue types (Supplementary file S8). These pathways synthesize precursors for various secondary metabolites and antioxidants vital for human health and thus make seeds more nutritious.

The correlation analysis gave us a hint of a significant relationship between highly upregulated transcription factors, and the other differentially expressed transcripts. We observed that MYB and C3H zinc finger transcription factors were highly upregulated in D69-Internode. Recent studies revealed that both transcription factor types are involved in internode elongation and development processes (Zhong et al., 2008; Kebrom, McKinley & Mullet, 2017; Gómez-Ariza et al., 2019). Sh_Salba_v2_112851, an AP2/ERF family member, is highly expressed in seed only and might play a role in dehydration-induced response as DREB2A proteins that are involved in response to drought, salt, and low-temperature stress (Nakashima et al., 2000; Sakuma et al., 2002). A set of 15 transcripts, correlated with Sh_Salba_v2_112851, were downregulated in seed, and participate in pathways that are downregulated in seed. For example-sh_salba_v2_ 33433 (CONSTANS-like 10) might be involved in the regulation of flowering genes (Tan et al., 2016), sh_salba_v2_01428 (histidine kinase 4) in cytokinin signaling (Ueguchi et al., 2001; Nishimura et al., 2004), sh_salba_v2_107585 (microtubule regulatory protein) in hypocotyl cell elongation (Liu et al., 2013), and sh_salba_v2_52914 (Apyrase) in normal growth and development of plant (Wolf et al., 2007). The correlation analysis suggests that transcription factors upregulated in seed and D69-internode tissues regulate various biological processes by controlling the expression of their target transcripts.

Further analysis of *de novo* assembled Chia transcriptome revealed 2,411 SSRs (see Supplementary file S11). Simple Sequence Repeats (SSRs) are an important class of genetic markers widely used in molecular breeding applications. SSRs identified in chia reference transcriptome might be a valuable resource for breeding and genetic improvement of the crop. Overall, this is the first study that generated a tissue-specific reference transcriptome atlas for a Chia, a neo model and an agronomically important crop.

## Materials and Methods

### Plant material, growth conditions and sampling

Seeds of Chia (*Salvia hispanica* L.) bought online from Ancient Naturals, LLC, Salba Corp, N.A. were sown in autoclaved soils and watered thoroughly under controlled greenhouse conditions. All seeds germinated on the third day after sowing. Since the primary seed material was expected to a heterogeneous mixture, biological replicates for each tissue type were collected from three randomly chosen plants. The description of the samples collected from various developmental stages and tissue types is shown in Table 1. The tissue samples include seeds, cotyledons, shoots from 3 and 12 days old seedlings, leaves from 12 (D12-P1) and 69 days old plants (D69-P1-P7), internode from 69 days old plants, raceme inflorescence from 158 days old plants, and flowers from 1 and 5 days post-anthesis. Collected samples were immediately frozen in liquid nitrogen and stored at −80°C.

### Sample preparation and sequencing

Total RNA from frozen tissues was extracted as per manufacturer’s protocol using RNA Plant reagent (Invitrogen Inc., USA), RNeasy kits (Qiagen Inc., USA), and treated with RNase-free DNase (Life Technologies Inc., USA). Total RNA concentration and quality were determined using ND-1000 spectrophotometer (Thermo Fisher Scientific Inc., USA) and Bioanalyzer 2100 (Agilent Technologies Inc., USA). Samples were prepared separately from each of the three biological replicates of each tissue type using the TruSeqTM RNA Sample Preparation Kits (v2) and sequenced using the Illumina HiSeq 2500 instrument (Illumina Inc., USA) at the Center for Genomic Research and Biocomputing, Oregon State University.

### De novo transcriptome assembly and quality estimation

FASTQ file generation from the RNA-Seq sequences was done by CASAVA software v1.8.2 (Illumina Inc.). Read quality was assessed using FastQC, and poor-quality reads were removed with Sickle v. 1.33 (-q = 20) (“najoshi/sickle”). The transcripts were assembled using Velvet (v1.2.10), which uses De Bruijn graphs to assemble short reads (Zerbino & Birney, 2008). An assembly of 67 and 71 k-mer lengths was performed using all tissue-specific reads. Assemblies produced by Velvet were merged into a single consensus assembly by Oases (v0.2.08) (Schulz et al., 2012), which produced transcript isoforms using read sequence and pairing information. Quality estimation to reduce redundancy in transcript assembly (a quality control check for *de novo* assembled transcriptome) was carried out using CD-HIT-EST (Li & Godzik, 2006), TransRate (Smith-Unna et al., 2016), and QUAST (Gurevich et al., 2013) software packages. The assembled transcripts passing the CD-HIT-EST quality control step were used for further downstream analyses and considered as a reference transcriptome for differential gene expression analyses.

### Functional annotation and pathway enrichment analysis

Assembled transcripts were annotated using BLASTx and tBLASTx with an E-value cutoff of 10^−10^. The assembled transcripts were translated into peptides using Transdecoder (v2.1.0) (“TransDecoder (Find Coding Regions Within Transcripts)”) with a minimum peptide length of 50 or more amino acids. Transdecoder used the BLASTp and PfamA search results to predict the translated ORF. Resulting peptides were analyzed using InterProScan Sequence Search (v5.17.56) (Zdobnov & Apweiler, 2001b; Jones et al., 2014b) hosted by the Discovery Environment and powered by CyVerse (Joyce et al., 2017). We used the AgriGO Analysis Toolkit (Du et al., 2010b) to identify statistically enriched function groups of transcripts. AgriGO uses a Fisher’s exact test with a Yekutieli correction for false discovery rate calculation. Significance cutoffs were set at a P-value of 0.05 and a minimum of 5 mapping entries per GO term. KAAS-KEGG automation server was used for orthologue assignment and pathway analysis (Moriya et al., 2007).

### Gene expression and clustering

Bowtie2 (Langmead & Salzberg, 2012) was used to align sequence reads from each tissue type to the assembled transcriptome. The RSEM software package (Li & Dewey, 2011a) was used to estimate the transcript expression counts (FPKM) from the aligned sequence reads. Count data obtained from RSEM was used in EBSeq (Leng et al., 2013) to identify differentially expressed genes based on the False Discovery Rate Corrected P-value of 0.05. Heatmaps were generated using Morpheus (Gould) developed by Broad Institute (https://software.broadinstitute.org/morpheus) and MEV (version 4.8.1) (*mev*, 2017) was used to cluster expression data from Chia. Log_2_ transformed fold change value for each transcript was used as input (p-value 0.1). Due to the orders of magnitude in the expression of transcripts between tissue types, we chose several methods of data normalization for cluster generation. Unit variance, median centering of transcripts, and summation of squares were applied to the dataset. In the investigation of individual gene families, transcripts were hierarchically clustered using a Pearson correlation.

### Coexpression and network analysis

The transcription factor transcripts were classified based on homology searches in Plant TFDB database v5.0 (http://planttfdb.cbi.pku.edu.cn) (Jin et al., 2017) and BlastX searches against *Arabidopsis thaliana*. For the coexpression analysis, CoExpNetViz tool (Tzfadia et al., 2015) was used. This tool utilizes a set of query or bait genes as an input and a gene expression dataset. Transcription factor transcripts displaying maximum expression cutoff of log_2_ transformed FPKM ≥ 5 were used as baits, and differentially expressed transcripts displaying maximum expression cutoff of log_2_ transformed FPKM ≥ 2 were used as expression matrix. Baits and expression matrix were loaded in CoExpNetViz tool, and the analysis was run to calculate coexpression with the setting of the Pearson correlation coefficient. For the expression matrix, transcripts considered as coexpressed if their correlation does not lie between the lower (5^th^) and upper (95^th^) percentile of the distribution of correlations between a sample of genes per gene expression matrix. The output files from the CoExpNetViz tool were used for displaying gene coexpression network using Cytoscape (version 3.7.1).

### Identification of Simple Sequence Repeats

Multiple length nucleotide SSRs were identified in the transcripts of the CD-HIT-EST assembly by using the stand-alone version of Simple Sequence Repeat Identification Tool (SSRIT) (Temnykh, 2001).

## Availability of supporting data

The raw sequencing data from all cDNA libraries were deposited at EMBL-EBI ArrayExpress (“ArrayExpress < EMBL-EBI”) under experiment number E-MTAB-5515.

## Supplementary Material

**Supplementary file S1:** A summary of the raw and clean reads obtained after the sequencing and preprocessing, respectively, and reads aligned to the reference transcriptome.

**Supplementary file S2:** Quality assessment of merged (column 2), CD-HIT-EST (column 3) and TransRate (column 4) assemblies using QUAST

**Supplementary file S3:** Workflow of Chia transcriptome sequencing and downstream analysis

**Supplementary file S4:** Functional annotation of chia peptides using InterProScan

**Supplementary file S5:** Gene Ontology annotations of chia peptides

**Supplementary file S6:** k-means clustering of transcripts depicting tissue□specific gene expression across different developmental stages. The Y-axis in each cluster denotes the mean-centered log_2_ transformed FPKM values ranging from +17 to −17.

**Supplementary file S7:** Transcripts clustered in 20 clusters

**Supplementary file S8:** Transcripts mapped to KEGG pathways

**Supplementary file S9:** Differentially expressed transcription factors across various developmental stages

**Supplementary file S10:** Frequency distribution of SSRs types in chia transcripts

**Supplementary file S11:** SSR motifs in chia transcripts

**Supplementary file S12:** SSRs identified in transcripts involved in metabolic pathways

