## Supplementary file S1 for "Chia (*Salvia hispanica*) gene expression atlas elucidates dynamic spatio-temporal changes associated with plant growth and development"

| **Sample name** | **Raw reads** | **High quality reads** | **Reads aligned** | **% reads aligned** |
| --- | --- | --- | --- | --- |
| Seed-Rep1 | 7,897,227 | 7,107,242 | 3,963,148 | 55.76 |
| Seed-Rep2 | 12,378,800 | 11,015,771 | 5,722,374 | 51.94 |
| Seed-Rep3 | 9,322,319 | 8,377,404 | 4,758,145 | 56.79 |
| D3-Cotyledon-Rep1 | 11,881,469 | 10,628,290 | 4,823,770 | 45.39 |
| D3-Cotyledon-Rep2 | 12,268,618 | 10,985,735 | 5,442,098 | 49.53 |
| D3-Cotyledon-Rep3 | 9,208,067 | 8,391,922 | 3,762,587 | 44.83 |
| D3-Shoot-Rep1 | 10,384,117 | 9,253,903 | 4,593,592 | 49.63 |
| D3-Shoot-Rep2 | 9,649,460 | 8,696,488 | 4,222,955 | 48.55 |
| D3-Shoot-Rep3 | 8,469,392 | 7,718,607 | 3,949,179 | 51.16 |
| D12-Shoot-Rep1 | 11,064,988 | 9,977,653 | 6,049,575 | 60.63 |
| D12-Shoot-Rep2 | 10,432,261 | 9,422,688 | 5,216,676 | 55.36 |
| D12-Shoot-Rep3 | 10,529,884 | 9,532,058 | 5,229,276 | 54.85 |
| D12-P1-Rep1 | 11,882,355 | 10,755,886 | 6,282,133 | 58.40 |
| D12-P1-Rep2 | 9,232,739 | 8,313,608 | 4,841,771 | 58.23 |
| D12-P1-Rep3 | 10,525,503 | 9,412,483 | 5,221,500 | 55.47 |
| D69-P1-P2-Rep1 | 11,128,184 | 10,066,319 | 6,315,653 | 62.74 |
| D69-P1-P2-Rep2 | 14,894,117 | 13,445,966 | 7,979,113 | 59.34 |
| D69-P1-P2-Rep3 | 8,513,792 | 7,722,686 | 4,709,083 | 60.97 |
| D69-P3-P4-Rep1 | 9,662,967 | 8,715,848 | 5,434,140 | 62.34 |
| D69-P3-P4-Rep2 | 10,187,987 | 9,196,112 | 5,092,594 | 55.37 |
| D69-P3-P4-Rep3 | 9,827,473 | 8,856,225 | 5,010,769 | 56.57 |
| D69-P5-P6-P7-Rep1 | 10,195,885 | 9,319,545 | 5,797,433 | 62.20 |
| D69-P5-P6-P7-Rep2 | 11,122,426 | 10,018,937 | 5,817,512 | 58.06 |
| D69-P5-P6-P7-Rep3 | 9,368,820 | 8,497,772 | 4,999,813 | 58.83 |
| D69-Internode-Rep1 | 9,480,054 | 8,469,710 | 5,135,626 | 60.63 |
| D69-Internode-Rep2 | 9,338,910 | 8,391,566 | 5,124,181 | 61.06 |
| D69-Internode-Rep3 | 7,894,588 | 7,083,391 | 5,042,654 | 71.18 |
| D158-RacemeTopHalf-Rep1 | 11,213,881 | 10,083,924 | 6,292,370 | 62.40 |
| D158-RacemeTopHalf-Rep2 | 10,726,327 | 9,672,346 | 5,826,664 | 60.24 |
| D158-RacemeTopHalf-Rep3 | 9,758,886 | 8,845,832 | 5,318,103 | 60.11 |
| D158-RacemeBottomHalf-Rep1 | 9,479,762 | 8,470,167 | 5,123,803 | 60.49 |
| D158-RacemeBottomHalf-Rep2 | 9,614,174 | 8,609,917 | 5,010,815 | 58.19 |
| D158-RacemeBottomHalf-Rep3 | 8,268,661 | 7,459,180 | 5,386,453 | 72.21 |
| D159-Flowers-Rep1 | 10,432,896 | 9,356,932 | 5,441,409 | 58.15 |
| D159-Flowers-Rep2 | 9,838,906 | 8,811,058 | 5,324,046 | 60.42 |
| D159-Flowers-Rep3 | 8,596,562 | 7,700,426 | 5,505,219 | 71.49 |
| D164-Flowers-Rep1 | 7,652,311 | 6,839,337 | 4,106,616 | 60.04 |
| D164-Flowers-Rep2 | 10,933,271 | 9,777,147 | 5,917,050 | 60.51 |
| D164-Flowers-Rep3 | 10,387,737 | 9,350,561 | 5,282,744 | 56.49 |

**Supplementary file S1:** A summary of the raw and clean reads obtained after the sequencing and preprocessing, respectively, and reads aligned to the reference transcriptome.
