## Supplementary file S2 for "Chia (*Salvia hispanica*) gene expression atlas elucidates dynamic spatio-temporal changes associated with plant growth and development"

Worst Median Best

☑ Show heatmap

| Statistics without reference | Sh_Salba_cDNA_v2 | Sh_Salba_v2-cdhittest | Sh_Salba_cDNA_v3 |
| --- | --- | --- | --- |
| # contigs | 145 503 | 82 663 | 35 461 |
| # contigs (>= 0 bp) | 145 503 | 82 663 | 35 461 |
| # contigs (>= 1000 bp) | 77 998 | 43 476 | 13 267 |
| # contigs (>= 5000 bp) | 931 | 466 | 26 |
| # contigs (>= 10000 bp) | 31 | 15 | 1 |
| # contigs (>= 25000 bp) | 0 | 0 | 0 |
| # contigs (>= 50000 bp) | 0 | 0 | 0 |
| Largest contig | 12 132 | 12 132 | 11 275 |
| Total length | 192 175 181 | 106 958 716 | 34 097 680 |
| Total length (>= 0 bp) | 192 175 181 | 106 958 716 | 34 097 680 |
| Total length (>= 1000 bp) | 153 862 642 | 85 176 225 | 22 349 024 |
| Total length (>= 5000 bp) | 5 618 432 | 2 800 052 | 154 260 |
| Total length (>= 10000 bp) | 334 870 | 163 142 | 11 275 |
| Total length (>= 25000 bp) | 0 | 0 | 0 |
| Total length (>= 50000 bp) | 0 | 0 | 0 |
| N50 | 1809 | 1791 | 1335 |
| N75 | 1146 | 1131 | 797 |
| L50 | 35 882 | 20 215 | 8685 |
| L75 | 68 921 | 38 821 | 16 870 |
| GC (%) | 40.54 | 40.07 | 41.2 |
| **Mismatches** | | | |
| # N's | 0 | 0 | 0 |
| # N's per 100 kbp | 0 | 0 | 0 |
