## Supplementary figures and images for "Chia (*Salvia hispanica*) gene expression atlas elucidates dynamic spatio-temporal changes associated with plant growth and development"

### Supplementary file S3

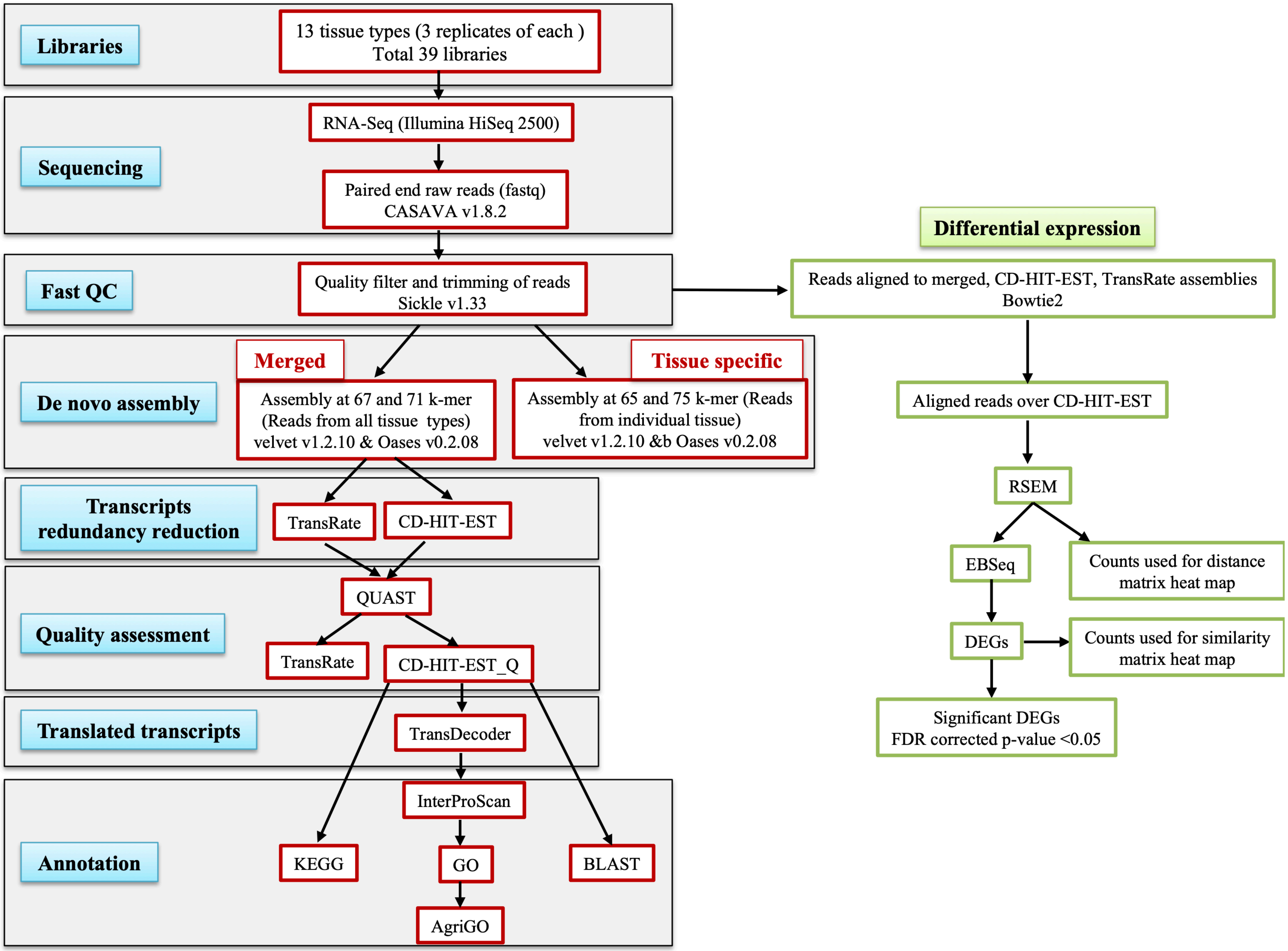

### Supplementary file S6

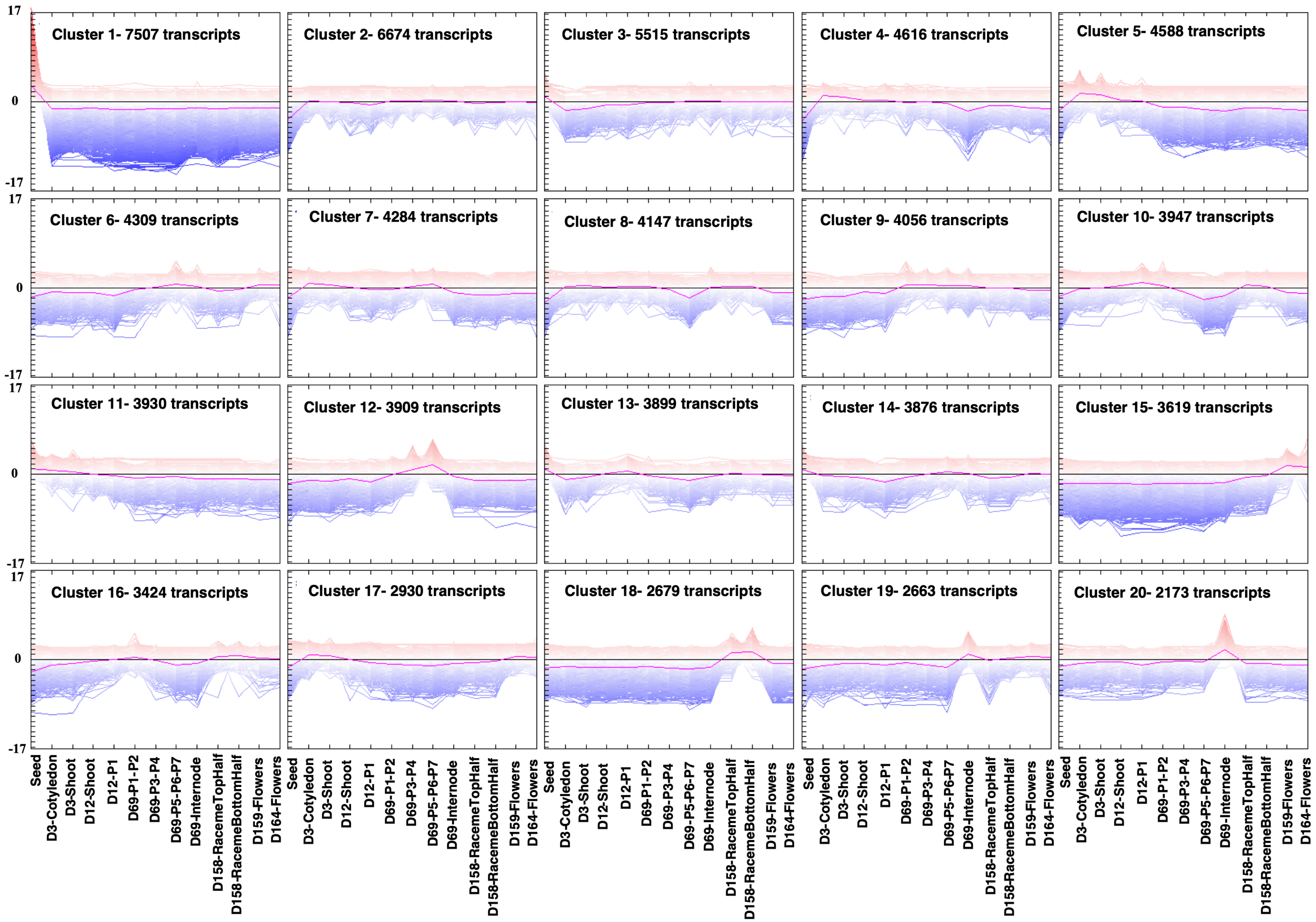

### Supplementary file S10

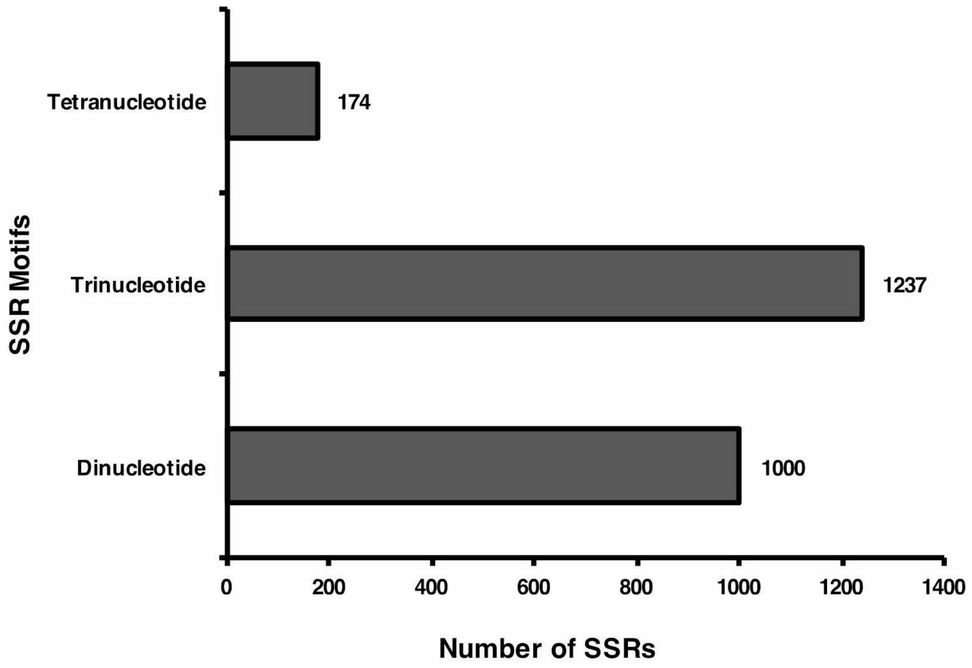
